# Post-GWAS functional analyses of *CNTNAP5* suggests its role in glaucomatous neurodegeneration

**DOI:** 10.1101/2024.03.14.583830

**Authors:** Sudipta Chakraborty, Jyotishman Sarma, Shantanu Saha Roy, Sukanya Mitra, Sayani Bagchi, Sankhadip Das, Sreemoyee Saha, Surajit Mahapatra, Samsiddhi Bhattacharjee, Mahua Maulik, Moulinath Acharya

## Abstract

Primary angle closure glaucoma (PACG) affects more than 20 million people worldwide, with an increased prevalence in south-east Asia. In a prior haplotype-based GWAS, we identified a novel *CNTNAP5* genic region, significantly associated with PACG. In the current study, we have extended our perception of *CNTNAP5* involvement in glaucomatous neurodegeneration in a zebrafish model, through investigating phenotypic consequences pertinent to retinal degeneration upon knockdown of cntnap5 by translation-blocking morpholinos. While cntnap5 knockdown was successfully validated using an antibody, immunofluorescence followed by western blot analyses in cntnap5-morphant (MO) zebrafish revealed increased expression of acetylated tubulin indicative of perturbed cytoarchitecture of retinal layers. Moreover, significant loss of Nissl substance is observed in the neuro-retinal layers of cntnap5-MO zebrafish eye, indicating neurodegeneration. Additionally, in spontaneous movement behavioural analysis, cntnap5-MO zebrafish have a significantly lower average distance traversed in light phase compared to mismatch-controls, whereas no significant difference was observed in the dark phase, corroborating with vision loss in the cntnap5-MO zebrafish. This study provides the first direct functional evidence of a putative role of *CNTNAP5* in visual neurodegeneration.

## Introduction

Primary angle closure glaucoma (PACG) is a major blinding eye disease, affecting over 20 million patients globally, more prevalent in east and south-east Asia^1^. It is characterized by progressive closure of the anterior chamber angle, resulting in blocked drainage of aqueous humour and increased intraocular pressure, which ultimately damages the optic nerve ^2,3^. Eventually, loss of retinal layers and the degeneration of the optic nerve can result in irreversible vision loss, leading to blindness ^4^. The disease has a strong heritable component, suggesting genetic factors play an important role in its pathogenesis^5^.

A few GWAS in Asian populations revealed associations between PACG and genetic variants in regions near or within the genes *PLEKHA7*, *COL11A1*, and *PCMTD1-ST18*^6^. Later, 5 new loci have been identified to be associated with disease risk near the genes *DPM2-FAM102A*, *FERMT2*, *GLIS3* and *CHAT*^7^. Notably, all these genes have known roles in ocular development, suggesting shared genetic mechanisms in angle development and closure. However, genomic factors contributing towards visual neurodegeneration aspect of the disease remains unexplored and elusive.

Previously, our extreme phenotype haplotype-based genome-wide association study identified *CNTNAP5* genic region as novel genomic loci to be significantly associated with PACG and one of its major endophenotype, cup-to-disc ratio (CDR)^8^. CDR is a clinical hallmark for glaucomatous neurodegeneration. Subsequently, our *in silico* analytical data further provided additional lines of evidence suggesting that *CNTNAP5* is implicated in potential expression changes and altered regulatory gene networks in glaucomatous pathophysiology^8^. *CNTNAP5* (Contactin Associated Protein 5) is a neural transmembrane protein, mainly enriched in myelinated axons that belongs to the neurexin superfamily^9^. It is a class of cell adhesion molecule of Contactin family, found primarily in the brain, where they play critical roles in neurite formation, neuronal development, axonal domain organisation, and axonal guidance^10^. Their absence causes axon malformation and poor nerve transmission. Earlier reports detected possible connection between rare variants in *CNTNAP5* with neurodevelopmental disorders or neurological diseases^11^. However, functional evidence validating the putative role of *CNTNAP5* in glaucomatous neurodegeneration remains to be determined.

There are numerous examples in the literature of GWAS loci that fail to be corroborated in functional studies, underscoring the need for rigorous follow-up analyses^12^. Moreover, translating GWAS findings into biological insights remains challenging, as the majority of disease-associated variants map to non-coding regions with unclear functional effects^13^. In this study, we have extended the current understanding of the molecular contributions of *CNTNAP5* to glaucomatous neurodegeneration. First, in order to incorporate genomic annotation data and emphasize *CNTNAP5* in order to search for their enhancer function and retinal expression, we used post-GWAS prioritization. We performed post-GWAS prioritization to integrate genomic annotation data. Next, we used translation-blocking morpholinos to knock down *CNTNAP*5 gene in zebrafish to study its possible involvement in a functional consequence towards dysregulated retinal development. Subsequently, we performed immunofluorescence and western blot analyses using an acetylated tubulin antibody to observe the expression status and retinal neural tissue architecture. Furthermore, we examined the effects of cntnap5 knockdown on apoptosis in zebrafish. In addition to immunofluorescence-based markers, we stained Nissl granules within neuronal cell bodies of retinal neurons to evaluate defects in ribosomal integrity reflective of neurodegeneration. Finally, in conjunction with the anatomical and histological analyses from genetic knockdown of cntnap5, we aimed at evaluating cntnap5 deficiency to precipitate measurable visual locomotory deficits equivalent to loss of vision in zebrafish.

## Materials and Methods

### Conditional analysis of *CNTNAP5* locus

To determine if the association signal for glaucomatous neurodegeneration previously identified through GWAS points to *CNTNAP5* as the likely causal gene within the chromosome 2 locus, we performed conditional and joint analysis using summary statistics of SNP genotype-phenotype associations across the region. This was carried out using version 1.93 of genome-wide complex trait analysis (GCTA) software^14^. LD-independent genome-wide significant SNPs within ±500kb window flanking *CNTNAP5* transcription start site were selected from the broader chromosome 2 locus showing evidence of association in the primary GWAS discovery analysis. The GCTA stepwise model selection (--cojo-slct algorithm) was applied conditioning on the top sentinel SNP across iterations to refine association signals, pinpoint candidate independent causal variants, and infer local genes mapping to said variants statistically accounting for most of the GWAS signal.

### Analysis of Cis-Regulatory Elements (CRE) annotations at *CNTNAP5* locus

To investigate the non-coding regulatory landscape surrounding prioritized candidate gene *CNTNAP5* for follow-up functional study, we searched for annotated CREs within the broader ±500 kb flanking region using the SCREEN DNA Elements Explorer platform^15^. We specifically queried genomic coordinates chr2:125,083,095-125,084,384 spanning the *CNTNAP5* locus for reported CREs characterized by empirical DNaseI, CTCF, H3K4me3, and H3K27ac signals. Data on genomic location, biochemical epigenetic readouts, normal cell/tissue source identifiers and other annotation metadata were systematically compiled for downstream analysis in R software (R, version 4.1.2) environment as described in Chakraborty et al, 2023^16^.

### Analysis of *CNTNAP5* SNPs in DeepSEA database and RegulomeDB

We input genomic coordinates and reference/alternate alleles for the 13 *CNTNAP5* intronic and 3’ UTR SNPs into the DeepSEA web server (http://deepsea.princeton.edu). Regulatory impacts of SNPs disrupting or creating binding sites and composite functional significance scores were compiled from DeepSEA multi-task predictions Exploring known and predicted regulatory potential of PACG-associated non-coding variants annotated to the *CNTNAP5* gene provides additional support linking the genetic association signals observed through orthogonal bioinformatics evidence^17^. To assess potential genetic regulation of *CNTNAP5*, we examined 13 identified SNPs in RegulomeDB. This database integrates various regulatory information including expression quantitative trait loci (eQTLs) and chromatin immunoprecipitation (ChIP-seq) data to predict regulatory potential of variants. SNPs were input and the resulting RegulomeDB scores were recorded, with lower scores indicating increased evidence for regulatory function.

### Analysis of ocular tissue expression patterns of cntnap5

To evaluate the expression patterns of *cntnap5* across ocular tissue types, we obtained RNA sequencing datasets from the eyeIntegration v2.12 (https://eyeintegration.nei.nih.gov/) and human eye transcriptomic atlas(https://www.eye-transcriptome.com/) encompassing tissues from the cornea, iris, lens, ciliary body, retinal pigment epithelium, choroid, central and peripheral retina.

### Exploring chromatin topology at *CNTNAP5* locus using 3D, 3DIV and HUGIn genome browser

To investigate potential long-range chromatin interactions connecting distal regulatory elements to prioritized candidate gene *CNTNAP5*, we utilized the 3D Genome Browser (http://3dgenome.org). We searched 3D Genome Browser encompassing the full *CNTNAP5* gene body along with ±500kb flanking regions. Interactive conformation plots centred on the *CNTNAP5* TSS were generated to identify significant long-range intra-TAD chromatin loop anchors that may harbour distal regulatory elements. By utilizing overlays with tissue-specific epigenomic annotations, this investigation of 3D genomic architecture maps surrounding *CNTNAP5* facilitates the identification of potential non-coding SNPs identified by GWAS signals that may have an impact on cntnap5 expression^18^. Additionally, we queried the same 13 SNPs of *CNTNAP5* in the 3DIV database^19^ to retrieve a list of statistically significant distal chromatin interaction partners and their genomic annotations. The distance normalized interaction frequencies and bias removed interaction frequencies were noted for each SNP. Further, we utilized the HUGIn browser^20^ to visualize and compare the observed versus expected read counts between the *CNTNAP5* gene locus and 13 SNPs of interest with distal chromatin regions in specifically in in the dorsolateral prefrontal cortex (DPLC) and neuronal progenitor cells (NPC). The raw observed and expected read counts were extracted for each SNP for descriptive statistical analysis. By leveraging these publicly available Hi-C data resources, we obtained processed long-range chromatin interaction data between our loci of interest and distal regulatory elements across a human neural cell and tissue types. The normalized interaction frequencies and raw read counts were utilized to characterize and compare the chromatin architecture surrounding the *CNTNAP5* and its 13 SNPs under study. **Gene Co-expression Analysis**

To identify genes co-expressed with *CNTNAP5*, we utilized two online databases - STRING (https://string-db.org/) and GeneMANIA (https://genemania.org/). In STRING, we input *CNTNAP5* and selected Homo sapiens as the organism. The analysis was run with a high confidence level (0.700). In GeneMANIA, we similarly input *CNTNAP5* with Homo sapiens selected. The co-expression analysis was run with automatic weighting. The resulting list of co-expressed genes from both databases were compiled.

### Pathway Enrichment Analysis

The combined list of *CNTNAP5* co-expressed genes was input into Enrichr^21^ for pathway enrichment analysis. The analysis was run against the KEGG and Reactome pathway databases with the default settings. Significantly enriched pathways were identified using an adjusted p-value cutoff of 0.05.

### Analysis of *CNTNAP5* variant-phenotype associations using HumanBase

To examine *CNTNAP5*, the gene was searched in the HumanBase database(https://hb.flatironinstitute.org/). The algorithm returns confidence scores predicting the likelihood of association between the queried gene and various diseases. These confidence scores are derived from the tissue-specific gene networks and analysis of known disease ontology terms.

### Analysis of *CNTNAP5* variant-phenotype associations using PheWAS

We queried the database of Phenotype-Wide Association Studies (PheWAS) catalog (phewascatalog.org). This aggregator tool combines published genotype-phenotype association statistics across a diversity of disease states profiled in large genomic datasets like the UK Biobank. We searched for all single nucleotide polymorphisms (SNPs) and small insertions/deletions mapped to within the genomic co-ordinates of *CNTNAP5* on chromosome 2 (chr2:125,083,095-125,084,384). Statistical association data linked to clinical diagnosis billing codes (ICD-10 classifications) were compiled across available PheWAS studies for variants catalogued within this *CNTNAP5* locus. Significantly associated phenotypes and disorders (at FDR or Bonferroni adjusted p-values < 0.05 threshold) were systematically tabulated and categorized by broad human diseases.

### Dual luciferase reporter assay

To determine our PACG associated prioritized SNP rs2553628 located downstream of the *CNTNAP5* locus exerts regulatory effects on enhancer activity; we carried out dual luciferase reporter assay. A 245 bp region flanking the rs2553628 variant was PCR amplified from PACG patient genomic DNA and directionally cloned into the multiple cloning site upstream of the firefly luciferase cassette using KpnI and XhoI restriction enzymes (NEB) in the pGL4.20 luciferase construct driven by an SV40 promoter (Promega). Patient and Controls subject’s DNA was used to generate the risk and reference allelic reporter constructs. HEK293 cells were cultured in DMEM (Gibco) supplemented with 10% FBS (Gibco) and seeded into 24-well plates at a density of 5x10^4^ cells per well. Cells were co-transfected after 24 hours with 50 ng of the CNTNAP5 luciferase reporter constructs along with 10 ng of pRL-TK Renilla luciferase internal control using FuGENE HD transfection reagent (Promega). At 48 hours post-transfection, cell lysates were collected and Firefly and Renilla luciferase signals measured using the Dual-Luciferase Reporter Assay System (Promega) as per manufacturer’s protocol on Tecan infinite 200 PRO. Normalized promoter activity was calculated as the ratio of Firefly to Renilla luminescence. Differences in normalized luciferase ratios between SNP alleles across three biological replicates were statistically compared by Student’s t-test.

### Zebrafish husbandry and maintenance

Zebrafish (*Danio rerio*) were maintained in recirculating Tecniplast benchtop rack systems under standard laboratory conditions on a 14-hour light/10-hour dark cycle. Adult fish were set up for natural spawning to obtain synchronized embryo clutches for experiments. Usually, zebrafish (Tubingen) begin breeding at first light (7 AM). After feeding at 6:30 PM on the preceding day, pairwise breeding was established on a sloped breeding tank filled with zebrafish water. In order to maintain a 2:1 female to male ratio, four females and two males were housed in the breeding tank. When the light came on at 7AM in the morning, they often breed the next day. Approximately 500 eggs were found at the base of the breeding tank. After utilizing a strainer to gather the eggs, the strainer was rinsed with 1X E3 media (5 mM NaCl, 0.17 mM KCl, 0.33 mM CaCl2, 0.33 mM MgSO4, and 10−5% methylene blue) before the eggs were put into the Petri dish (with an average of 25 embryos/Petri dish) and kept in the incubator at 28.4 °C. All experiments in this study were conducted in compliance with institutional animal protocols under Committee for Control and Supervision of Experiments on Animal (CPCSEA) approval (No. NIBMGZF/002/2022-23).

### Gross morphological eye measurements

At 96 hpf (hour post fertilization), larval zebrafish were anesthetized in tricaine solution and lateral images captured under a stereomicroscope (LEICA M205 FA) for morphological measurements. Eye diameter, defined as the axial length between the nasal and temporal periphery across the centre, was quantified for each larva using LAS 4V.13 software (LEICA). A 2-sample t-test was performed to compare the gross anatomical eye size both vertical and horizontal diameter of morphants and mismatch control in zebrafish.

### Microinjection of morpholino

Translation blocker (TB) morpholinos and 5 base mismatch morpholino against cntnap5a and cntnap5b were purchased from Gene Tools, LLC. MO sequences are listed in Supplementary Material, **Table S1**. MO stocks were diluted in 10 mM KCl and 25% Phenol Red and each morpholino (2 ng) were mixed in a volume of 2 nL was microinjected injected into 1–2 cell-stage embryos using calibrated injection volumes^22^.

### Tissue processing and cryosectioning

96 hpf zebrafish were anesthetized in tricaine methane sulfonate and subsequently euthanized following approved animal care protocols (Ferguson et al, 2019). Zebrafish were fixed overnight at 4°C in 4% paraformaldehyde (PFA) in phosphate buffered saline (PBS). After fixation, samples were washed in PBS 2-3 times to remove residual PFA. Tissues were then immersed in 30% sucrose in PBS overnight at 4°C for cryoprotection. Subsequently, zebrafish were oriented in OCT compound in peel-away cryomolds and quickly frozen using dry ice. Cryosections were cut at 15 μm thickness using a cryostat (make) (LEICA CM 1860 UV) and collected directly on charged slides (Autofrost, Cancer Diagnostics). Sections were adhered to the slides by drying overnight at room temperature and stored at -80°C until use. Prior to immunostaining, slides were thawed at room temperature for 30-60 minutes and dried completely before beginning staining procedures.

### Eosin and Hematoxylin staining

The sections on glass slides were stained as follows: PBS for 12, Mayer’s hematoxylin (SRL) in aqueous solution for 15 sec, water for 15 sec, eosin (SRL) in ethanolic solution for 20 sec, followed by dehydration with 100% ethanol for 5 sec twice, and finally mounted in DPX (Sigma) with cover slip. The tissue sections were observed under the light microscope (EVOS XL core Invitrogen).

### Immunofluorescence

Immunofluorescence staining on cryosections was performed following previously published protocols (Ferguson et al 2019) Briefly, Cryosections were thawed and dried at room temperature for 30 minutes prior to immunostaining. Tissues were rehydrated with PBS and blocked for 1 hour at room temperature with 5% normal goat serum ((Gibco) source) in PBS containing 0.3% Triton X-100. Sections were incubated with primary antibodies overnight at 4°C. The following primary antibodies and dilutions were used: mouse anti-acetylated tubulin (1:1000, T7451, Sigma Aldrich), rabbit anti-cntnap5 (1:200, Abgenex). After overnight primary antibody incubation, slides were washed 3 times with PBS and incubated with isotype-specific anti-rabbit secondary antibody labeled with Alexa Fluor 488 (1:500, Abcam) and anti-mouse secondary antibody labeled with Alexa Fluor 568 (1:500, Abcam). AlexaFluor-conjugated secondary antibodies (1:1000) for 2 hours at room temperature protected from light. The nuclei were stained with 4′,6-diamidino-2-phenylindole (DAPI) (1 μg/mL, Invitrogen) for 5 min at room temperature. Finally, sections were washed with 1X PBS and mounted with ProLong Gold Antifade Mountant (Invitrogen). Images were acquired using confocal microscopy (Nikon Ti2 Eclipse). Image analysis was conducted using Nikon NIS-Elements software and ROI intensity was compared by Student t-test using GraphPad Prism software (v9, Dormatics).

### Generation of customized cntnap5 antibody for gene knockdown validation

The sequence of cntnap5 Protein (XP_009294486.1 5 isoform X2, *Danio rerio*) was analysed by Hydroplotter software from protein lounge (proteinlounge.com) to select the suitable epitopes for antibody generation. The selection is based on analysing the amino acid sequences in Kyte-Doolittle and Hopp-Woods plots and selecting a suitable hydrophilic region also Kolaskar Tongaonkar for antigenicity. Based on the above analysis we have selected RSERNVREASLQVDQLPLR (571-589aa) for antibody development. (Antigenicity Index: 1.028, Hydrophobicity Index: -1.084, Hydrophilicity Index: 1.268)

A cysteine has been added in the N-term of the petite for conjugation to carrier molecule. After synthesis of the peptide, it is conjugated to KLH (Keyhole lymphatic hemocyanin).

Then it was immunized to the two healthy New Zealand white rabbits with freund’s complete adjuvant (CFA) followed by freund’s incomplete adjuvant (IFA). After four boosters the first immune sera were collected. The initial screening of the developed antisera was carried out by coating ELISA plate with the unconjugated peptide (200ng/well) (**Figure S1A)**.

Based on the above information Rabbit B 3rd bleed was selected for further peptide specific antibody purification by conjugating the free peptide with Sepharose 4B matrix. The purified antibody was again validated by employing indirect ELISA (**Figure S1B**).

### Whole Mount Immunofluorescence

At 32hpf, embryos were dechorinated and transferred in 4% 1x PBS. Samples were permeabilized in ice-cold 100% methanol drop wise at -20°C for 2 hour and then washed with 1x PDT. Non-specific binding was blocked with 5% goat serum, 1% bovine serum albumin in PBS-Tween for 1 hr at room temperature. Embryos were incubated overnight at 4°C with rabbit anti-active caspase-3 primary antibody (1:500; Cell Signalling Technology) to label apoptotic cells. On the following day, embryos were washed and incubated with Alexa Fluor 568 goat anti-rabbit IgG secondary antibody (1:500; Molecular Probes) for 2 hrs at room temperature. Samples were washed with 1x PDT prior to imaging. 4% methylcellulose was used for whole mount imaging under the confocal microscope^23^.

### NeuroTrace Staining

Cryosections ranging from 12-15 μm thickness were collected on charged slides. Zebrafish eye tissue sections were stained with NeuroTrace 435/455 Blue Fluorescent Nissl Stain (1:50, N21479, Thermo Fisher) for 30 mins at room temperature as per manufacturer’s instructions. Sections were then washed 3 times in PBS and mounted with ProLong Gold Antifade (Invitrogen). Images were captured in confocal microscopy (Nikon Ti2 Eclipse). Image intensity was measured using Nikon NIS-Elements software. The whole eye was selected as region of interest (ROI) and the fluorescence intensity within the designated ROIs was quantified. In order to account for non-specific fluorescence, background correction was done when recording mean intensity values. ROI intensity was compared by Student’s 2-sample t-test using GraphPad Prism software (v9, Dormatics).

### Zebrafish protein isolation

Zebrafish embryos were collected for protein isolation at 96 hpf after morpholino injection. Embryos were washed with PBS three times to remove any E3 media remaining. Whole cell lysis buffer was added to the embryos (HEPES (20mM) pH 7.6, 100mM NaCl, 1.5mM MgCl_2_, 1mM DTT, 0.1% Triton X-100, 20% Glycerol and freshly added 10ul 0.1 M PMSF, 5ul Protease inhibitor cocktail per mL of buffer) and homogenized using cordless mortar (Sigma-Aldrich, Z359971) and pestle (Sigma-Aldrich, Z359947) keeping in ice. After incubation in ice for 30 minutes the samples were centrifuged at 13,000 rpm for 20 minutes at 4°C. The supernatant was collected and used in western blotting^24^.

### Western Blotting

Protein concentration in the prepared zebrafish protein lysate were estimated using Bradford reagent (Bio-Rad). Equal amounts of protein (30ug) were resolved by 10% sodium dodecyl sulphate–polyacrylamide gel electrophoresis. proteins were transferred to nitrocellulose membrane. Membranes were blocked with 5% non-fat milk in TBST (Tris-buffered saline-Tween 20) for 1 hour and incubated with primary antibodies overnight at 4°C. The following primary and secondary antibodies were used as per the mentioned dilutions: anti Acetyl-α-Tubulin Antibody (T7451, Sigma Aldrich), anti GAPDH (1:1000, 2118, CST), Goat Anti-Mouse IgG H&L (1:10,000, ab6708, Abcam), Goat Anti-Rabbit IgG H&L (HRP) (1:10,000, ab6721, Abcam). The blots were visualized using SuperSignal™ West Pico PLUS Chemiluminescent Substrate (Thermofisher) in ChemiDoc XRS+ System (Bio-Rad). Densitometric quantification was measured in unsaturated images using ImageJ software (National Institutes of Health, Bethesda, MD, USA).

### Spontaneous movement analysis

Larval zebrafish at 96 hpf were arrayed into 6-well transparent microtiter plate (1 larvae per well and each time 3 well for mismatch controls and 3 well for morphants) containing 200 µL E3 medium in each well. Plates were loaded into the Zantiks MVP system for locomotor tracking. The system was calibrated to maintain constant 28°C temperature and stabilize lighting for 30 minutes prior to recording. Following a 60-minute habituation period under darkness, the integrated software was programmed for automated alternating 10-minute epochs of light on and light off with complete darkness over a total 60-minute period. The movement tracks and total distance travelled (mm) per well were quantified for each light/dark interval period by the Zantiks tracking software. Statistical analysis was conducted using GraphPad Prism software (v9, Dormatics). A nonparametric Mann-Whitney test was performed to compare the groups in light and dark separately.

## Results

### In silico and in vitro characterization of CNTNAP5 genic region

Conditional analysis of thirteen genome-wide significant SNPs from our previous study clustered within a ∼500kb region of *CNTNAP5* on chromosome 2 showed maintenance of statistically significant genomic association with PACG. This indicates *CNTNAP5* is likely a candidate gene for the glaucomatous trait (**Figure 1A**). Next, bokeh plot of linkage disequilibrium (LD) revealed that our earlier associated 13 variants of *CNTNAP5*, clustered into two LD block (**Figure 1B**). To contextualize the 13 PACG-associated intronic and downstream *CNTNAP5* variants, uncovered from conditional analysis in terms of their putative regulatory functions, we intersected their locations with candidate cis-regulatory elements (CREs) annotated in the region. Plotting the *CNTNAP5* SNPs relative to adjacent CRE genomic positions revealed two distinct spatial clusters exhibiting differences in predicted regulatory element abundance. One cluster (cluster I, Blue Square) containing rs17011381, rs2901264, rs2115890, rs733112, rs1430263, rs17011394 and rs780010) reside in a genomically sparse region largely devoid of annotated CREs. In contrast, the cluster II (Red square) encompassing rs17724018, rs2553625, rs17011420, rs2553628 and rs17011429 directly overlap with a dense hotspot of CREs enriched in putative active chromatin marks (**Figure 1C**). This differential CRE density profile provides preliminary evidence suggesting the latter SNP cluster has a higher probability of harboring non-coding variants interfering with functionally relevant gene regulatory elements that may contribute to a breakdown in *CNTNAP5* expression. Additionally, in DeepSEA database, we found same set (cluster II) of SNP are having significant enrichment scores on histone modifications demarcating putative active enhancer elements within introns of *CNTNAP*5 (**Figure 1D**). The RegulomeDB score of these 13 SNPs were enlisted in the **Table S2**. The analysis of regulatory features for SNPs rs2553628 and rs17011429 revealed that both variants have high gene regulatory potential and both are residing in a strong linkage disequilibrium (LD). Examination of the HaploReg database showed that rs2553628 alters the motif for transcription factor TCF4 and having a CTCF binding sites, while rs17011429 alters motifs for GR and Hsf. TCF4 is reportedly associated with neurological disorders^25^. Furthermore, rs2553628 demonstrated significantly altered motif activity between the reference and alternative alleles (Figure 2A). To functionally check the enhancer activity, in the dual luciferase assay, the G allele of rs2553628 (from the cluster II) was showing the significantly higher luciferase activity than the A allele (P value =0.0034) (**Figure 2B** and **Figure S2**). On further exploration genomic architectural aspects surrounding *CNTNAP5*, we visualized Hi-C chromosomal conformation contact matrices encompassing the gene locus using the 3D Genome Browser. Notably, *CNTNAP5* resides within a topologically associating domain (TAD) suggesting capacity for long-range spatial contacts between the gene body and more distal enhancer elements (**Figure S3A**). The contact map, as well as the corresponding arc also suggests distal as well as proximal interactions in the regions surrounding the prioritized genic region in the dorsolateral prefrontal cortex (**Figure S3B**). Moreover, the HUGIn analysis of the genomic region of *CNTNAP5* reveals high proximal as well as distal interactions in the surrounding area as the observed counts were more than the expected counts especially outside the gene. The 4C plot of database shows the presence of the frequently interacting regions (FIRE) near the *CNTNAP5* genic region in the DPLC dorsolateral prefrontal cortex (DPLC) peaks for H3K27ac, H3k4me1 and H3k4me3 peaks in both DPLC and neuronal progenitor cells (NPC) (**Figure 2 C-E and Table S3**). The Human Protein Atlas expression dataset showed that the expression of *CNTNAP5* is restricted is only in retinal and neural tissue **(Figure S4**). Analysis of human eye transcriptomic atlas datasets revealed marked enrichment of *CNTNAP5* expression within central retinal tissues amongst profiled ocular cell types (**Figure S5).** The neural pathways were significantly enriched after performing pathways enrichment analysis of the co-expressed genes of *CNTNAP5* (**Figure S6 and S7**). Analysis of the *CNTNAP5* gene in the HumanBase database revealed several disease associations. The strongest association was with autism spectrum disorder, with a confidence score of 0.54. A weaker association was also found with epilepsy (confidence score 0.11). Most notably, HumanBase analysis also predicted an association between *CNTNAP5* and retinal disease, with a confidence score of 0.11. To visualize these results, the confidence scores for *CNTNAP5* across various neurological and retinal diseases were shown (**Table S4**). Together, with the genetic and genomic architectural evidence, the expression patterns further strengthen the candidacy of *CNTNAP5* in contributing to retinal phenotype outcomes in PACG. Additionally, mining of the phenotype-wide association study (PheWAS) catalog revealed significant genetic associations between variants in the *CNTNAP5* locus and several neurological disorders including epilepsy, schizophrenia and Parkinson’s disease (**Figure S8**).

**Figure 1:**
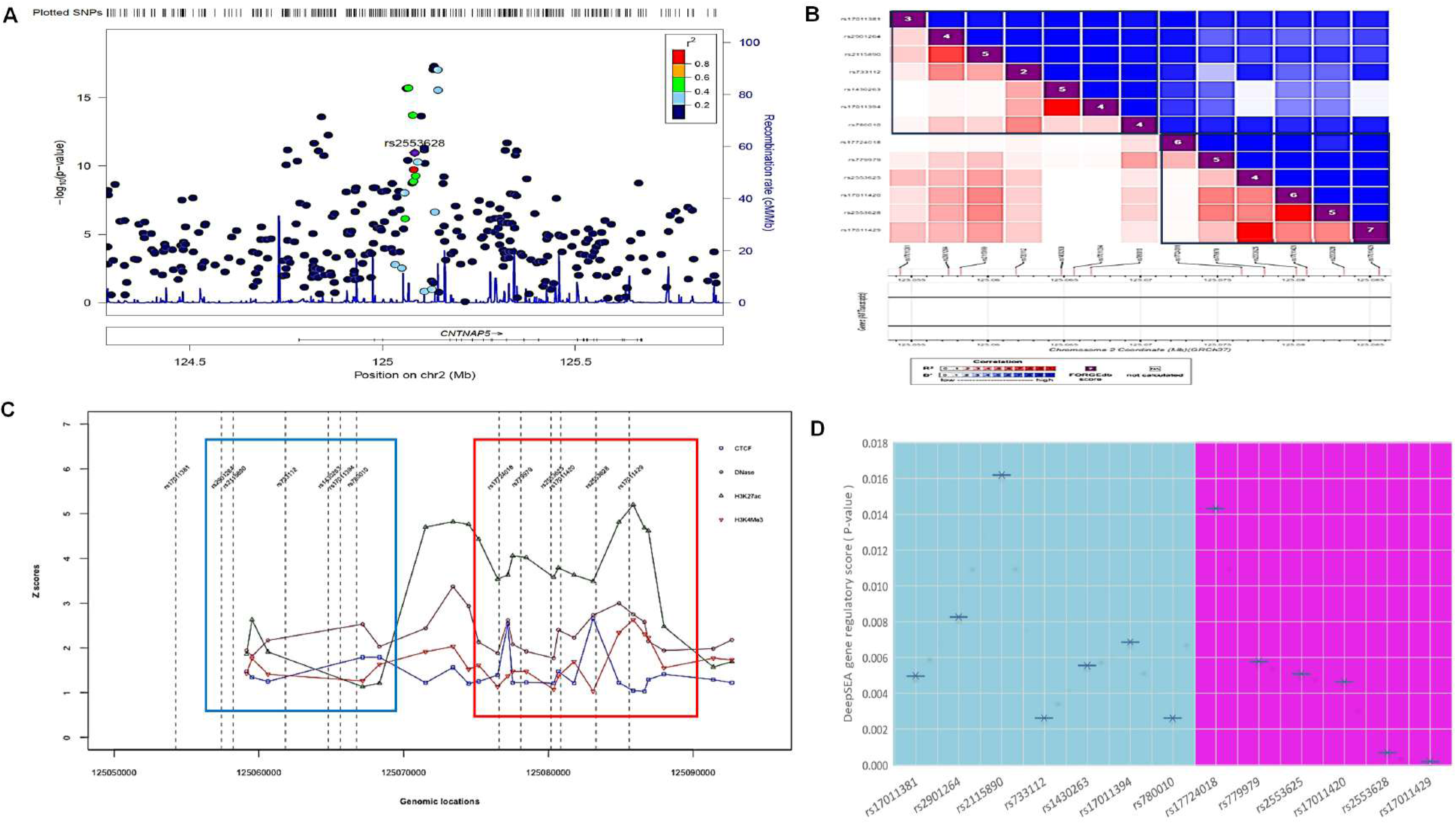
A. Regional Association Plot indicating the statistical strength of association among associated genomic region of *CNTNAP5* gene with PACG after performing conditional analysis **B.** Bokeh plot shows the pairwise linkage disequilibrium between the 13 variants around the region of *CNTNAP5* in two LD blocks **C.** CREs and SNPs plotted along with their genomic and epigenomic features of 13 SNPs of *CNTNAP5*. Based on the features, blue and red boxes are shown that roughly highlight SNP and CRE clusters. The elements and variants belonging to the red box show better likelihood for having functional relevance. **D.** The Whisker plot shows the DeepSEA gene regulatory score (P-value) of 13 SNPs of CNTNAP5. Based on the features, blue (cluster I) and pink (cluster II) boxes are shown that represents DeepSEA gene regulatory score (P-value). The elements and variants belonging to the pink square better likelihood for having gene regulation.

**Figure 2:**
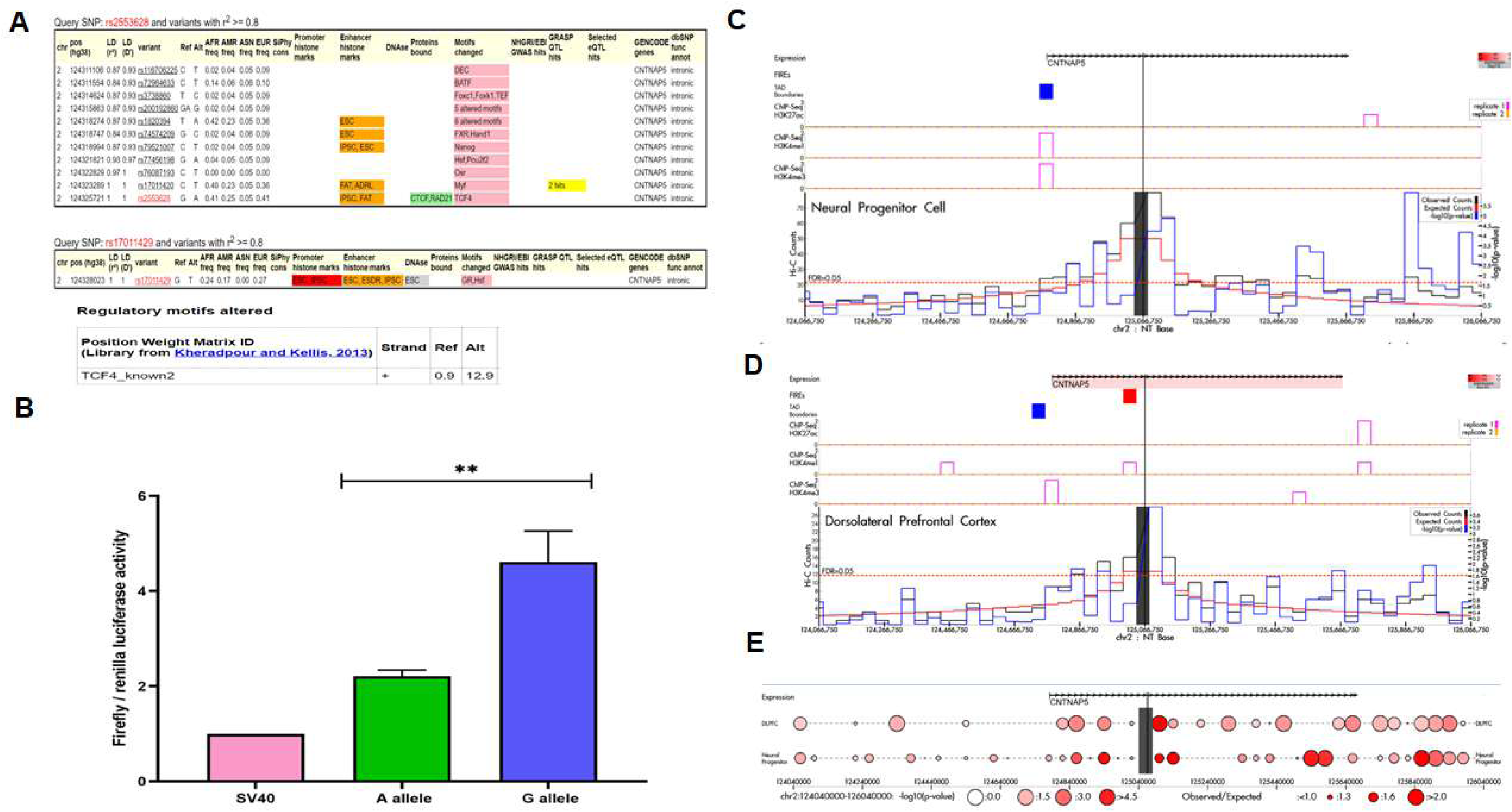
A. Variant detailed view of rs2553658 and rs17011429 in HaploReg. The regulatory motif TCF4 altered in reference and alternative allele of rs2553658. **B.** The data was drawn for a total of 3 biological replicates considering triplicate technical replicates for each biological experiments representing three different plasmid preparations (pGL3 SV40 minimal promoter vector) transfected in to HEK293T cells grown in 96-well plates. The height of the bar shows the mean value of all the 3 experiments, while error bars are depicting SE. The 4C plot of genomic region of CNTNAP5 (chr2:125,083,095-125,084,384) showing the FIREs, ChIP-Seq data and Hi-C counts in DPLC **(C)** and NPC **(D)** from HUGIn. The ratio of observed read counts to expected read counts is higher in NPC and is also statistically more significant than in DPLC **(E).**

*Cntnap5 morpholino-injected embryos show ocular phenotypic alterations in zebrafish* Following extensive genetic fine-mapping data integration indicating *CNTNAP5* as a putative contributor in glaucomatous neurodegeneration, we next sought direct functional validation *in vivo* using antisense knockdown approaches in zebrafish larvae. Quantitative gross anatomical phenotyping revealed significantly reduced diameters along both the nasal-temporal axis (horizontal) and superior-inferior axis (vertical) of eyes measured from cntnap5 a and b translation-blocking double morpholino (dMO) injected embryos (n=72, biological replicates = 6) at 72 hours post fertilization compared to 5 base pair mismatch (MM) morpholino controls (n=72, biological replicates = 6) (P<0.001) (**Figure 3A and B**). There was no significant death observed in both control and morphant groups. The injection rate, death rate and other anatomical parameter was compared in both groups (**Figure S9**). In deep phenotyping, histological examination showed noticeably diminished retinal nuclear layers with gaps and discontinuities in laminar organization in haematoxylin & eosin-stained thin sections from cntnap5 deficient larvae (**Figure 3C**).

**Figure 3:**
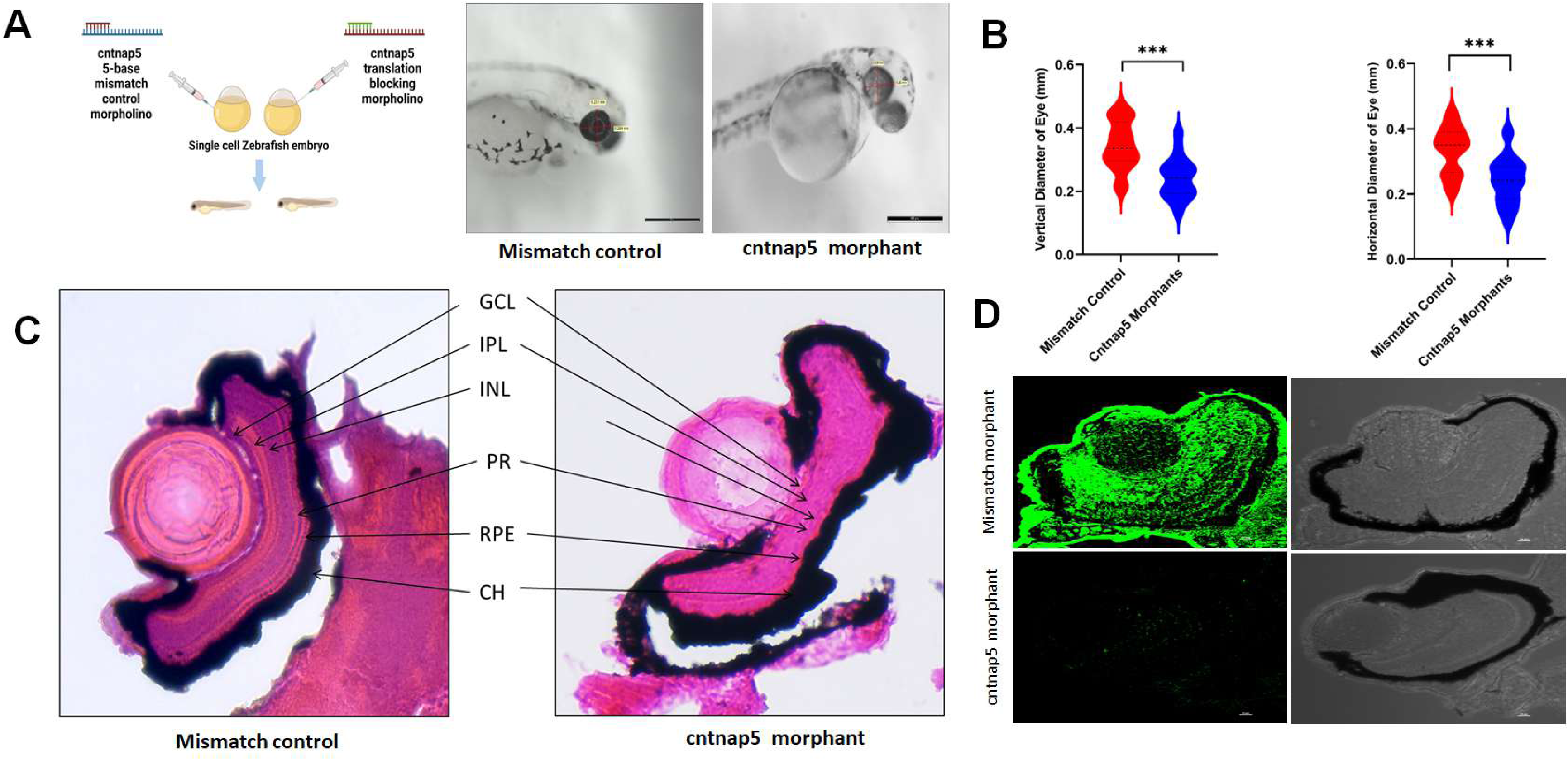
A. Morpholino knockdown of *CNTNAP5*. Zebrafish were microinjected with a cntnap5 translation blocking morpholino. Images taken 96 hpf, eye of a mismatch morphant (left) and an eye cntnap5 morpholino injected zebrafish (right) showing smaller anatomical eye than mismatch control**. B.** Quantitative gross anatomical diameters along both the nasal-temporal axis (horizontal) and dorsal-ventral axis (vertical) of eyes measured from cntnap5 translation-blocking morpholino injected embryos and mismatch morpholino controls zebrafish at 96 hours post fertilization stage. bars = mean ± SD, ns not significant, \*\*\**p*L<L0.005. **C.** Histological analysis of zebrafish eye microinjected with mismatch morpholino (left) a cntnap5 translation blocking morpholino (right) using H&E stain at 96 hpf stage showing disturbed retinal layers in cntnap5 morphant. GCL, ganglion cell layer; IPL, inner plexiform layer; INL, inner nuclear layer; PR, photoreceptor layer; RPE, retinal pigment epithelium; CH indicates choroid. **D.** Representative IF image of tissues stained with anti-cntnap5 showing cntnap5 expression in mismatch morphant (upper left; lower Transmitted Light Differential Interference Contrast (TD) image) but no expression in a cntnap5 translation blocking morphants (lower left: right: TD image).

### Cntnap5 antibody mediated validation of gene knockdown

In conjunction with the gross morphological and histological analyses demonstrating perturbation of eye and retinal development upon repression of endogenous cntnap5 translation, we also aimed to directly confirm successful protein knockdown in our zebrafish model using a target-specific customized antibody. As expected, dMOs exhibited a near complete absence of detectable anti-cntnap5 immunofluorescence throughout the retinal layers of eye. In contrast, the mismatch controls clear cntnap5 antibody immunostaining in their whole retinal layers, P value <0.0005 (**Figure 3D** and **Figure S10**). Thus, our data clearly suggest that cntnap5 deficiency disrupts retinal development in larval zebrafish.

### Cntnap5 morpholino-injected embryo eye showed disturbed retinal cytoarchitecture

To investigate the retinal tissue architecture after deep phenotyping using H&E staining, remarkably, the morphant retina showed profoundly increased anti-acetylated tubulin immunofluorescence signal intensity and disoriented retinal tissue organization throughout the retinal ganglion cell layer compared to mismatch morphant retina; P value = 0.0108 (**Figure S11**). To validate this, we co-stained with anti-cntnpa5 antibody along with DAPI to stain the nuclei (**Figure 4A**). Similarly, western blot quantification of whole embryo lysates further corroborated a significant ∼1.5-fold upregulation of acetylated alpha tubulin protein levels in cntnap5 deficient embryos relative to mismatch controls; P value = 0.031 (**Figure 4B**).

**Figure 4:**
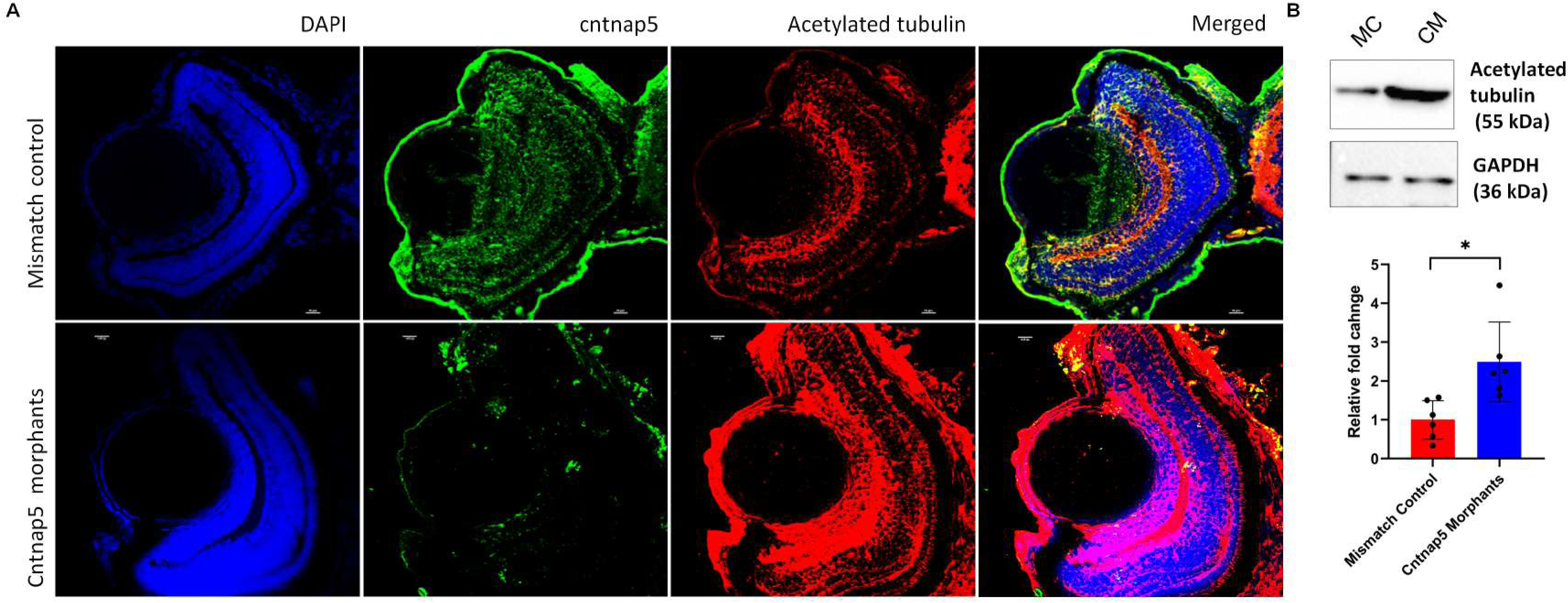
A. Representative confocal images of cntnap5 and acetylated tubulin expression of eye tissues from mismatch control fish (upper) and cntnap5 morphant (lower) zebrafish at 96 hpf (blue-DAPI, green-cntnap5, red-acetylated tubulin, and merged)**. B.** Western blot image of acetylated tubulin expression of whole zebrafish tissue lysate (72 hpf). MO: cntnap5 morpholino injected fish MM: mismatch control morpholino injected fish. A two-sided t-test. bars = mean ± SD, ns not significant, \**p*L<L0.05.

### Cntnap5 morpholino-injected zebrafish embryos show visual neurodegeneration and greater apoptosis

During our investigation, we employed the NeuroTrace stain to examine zebrafish retinal neurons. Notably, our findings revealed a significant reduction in NeuroTrace stain mirroring less Nissil granule intensity of retinal neurons within the dMOs compared to the MMs (P<0.005) (**Figure 5A** & **5B**). This observed decrease in NeuroTrace stain intensity strongly suggests the presence of neurodegeneration in the cntnap5 morphant zebrafish. Finally, we performed to observe the apoptosis by cleaved caspase 3 as an executioner caspase at a system level. Whole mount cleaved caspase 3 immunofluorescence staining showed dMOs had significantly more cleaved caspase 3 staining compared to mismatch controls (P<0.005). This indicates greater apoptosis in the dMOs versus controls (**Figure 5C** & **5D**). Together, these results demonstrate increased apoptosis in vivo in dMOs, supporting the importance of cntnap5 for cell survival.

**Figure 5:**
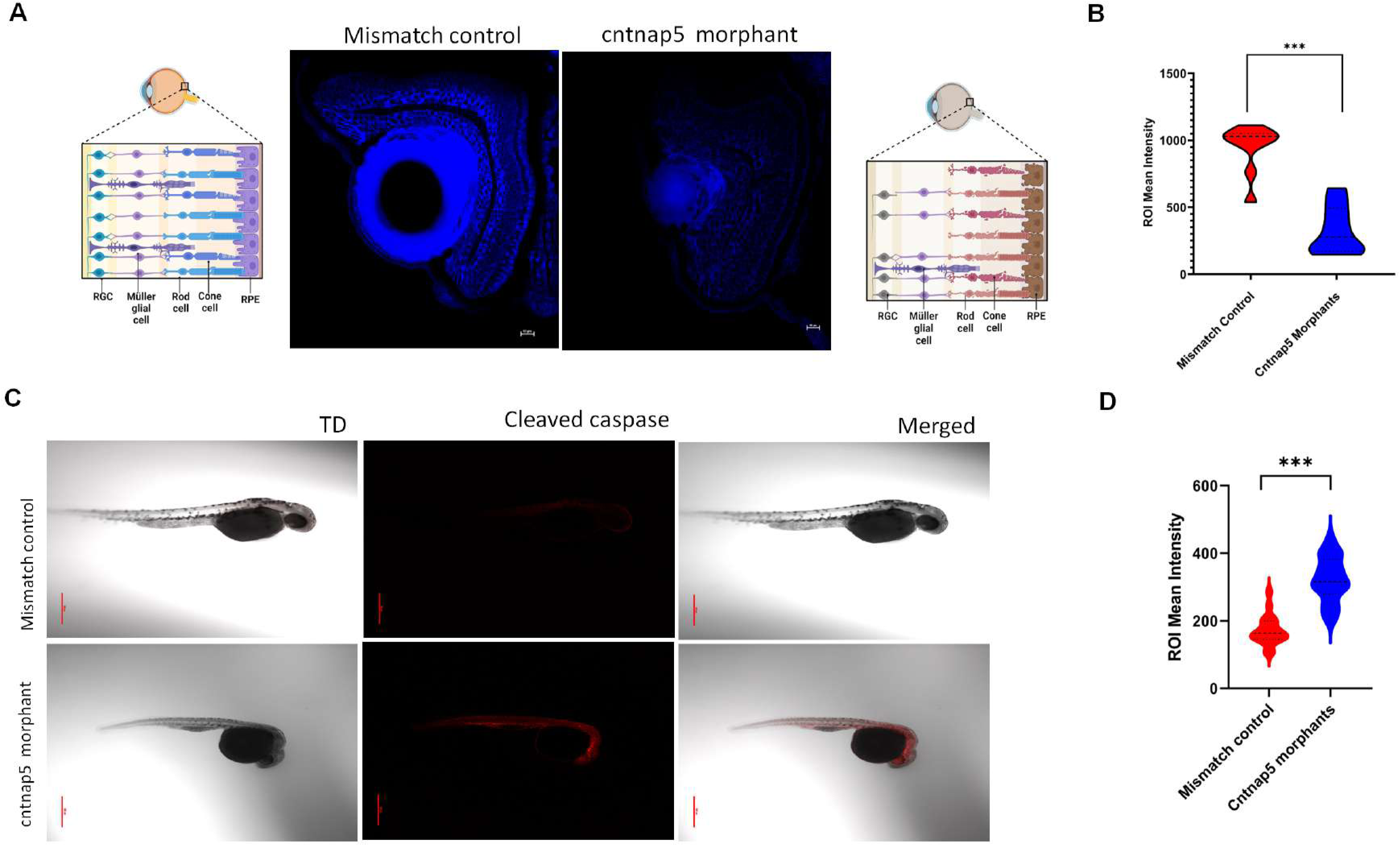
A. Representative eye tissue images of Nissl granules expression of zebrafish embryos (96 hpf) injected with a mismatch control (left) and cntnap5 morpholino (right). **B.** Comparative analysis of eye mean intensity of eye for both groups. bars = mean ± SE, ns not significant, \*\*\**p*L<L0.005. **C.** Representative whole mount confocal images of cleaved expression of zebrafish embryos (32 hpf) injected with cntnap5 morpholino (upper) or mismatch control (lower) **D.** Comparative analysis of mean intensity of eye for the both groups, bars = mean ± SE, ns not significant, \*\*\**p*L<L0.005.

### Spontaneous locomotory movement in cntnap5 knockdown fish in light and dark phase

We analyzed the effects of cntnap5 knockdown on visually guided locomotor activity under intermittent light/dark conditions using the Zantiks system. During 10-minute light periods, cntnap5 morphants (dMO) showed significantly less average swimming distance compared to mismatch control larvae (p<0.005), reflecting impaired significant locomotory movement in bright light phase. However, distance moved in dark conditions were not significantly different between groups (p=0.23) (**Figure 6A**). In summary, cntnap5 morphant zebrafish spontaneous movement was impaired in light phase compared to mismatch controls indicating their vision loss (**Figure 6B**).

**Figure 6:**
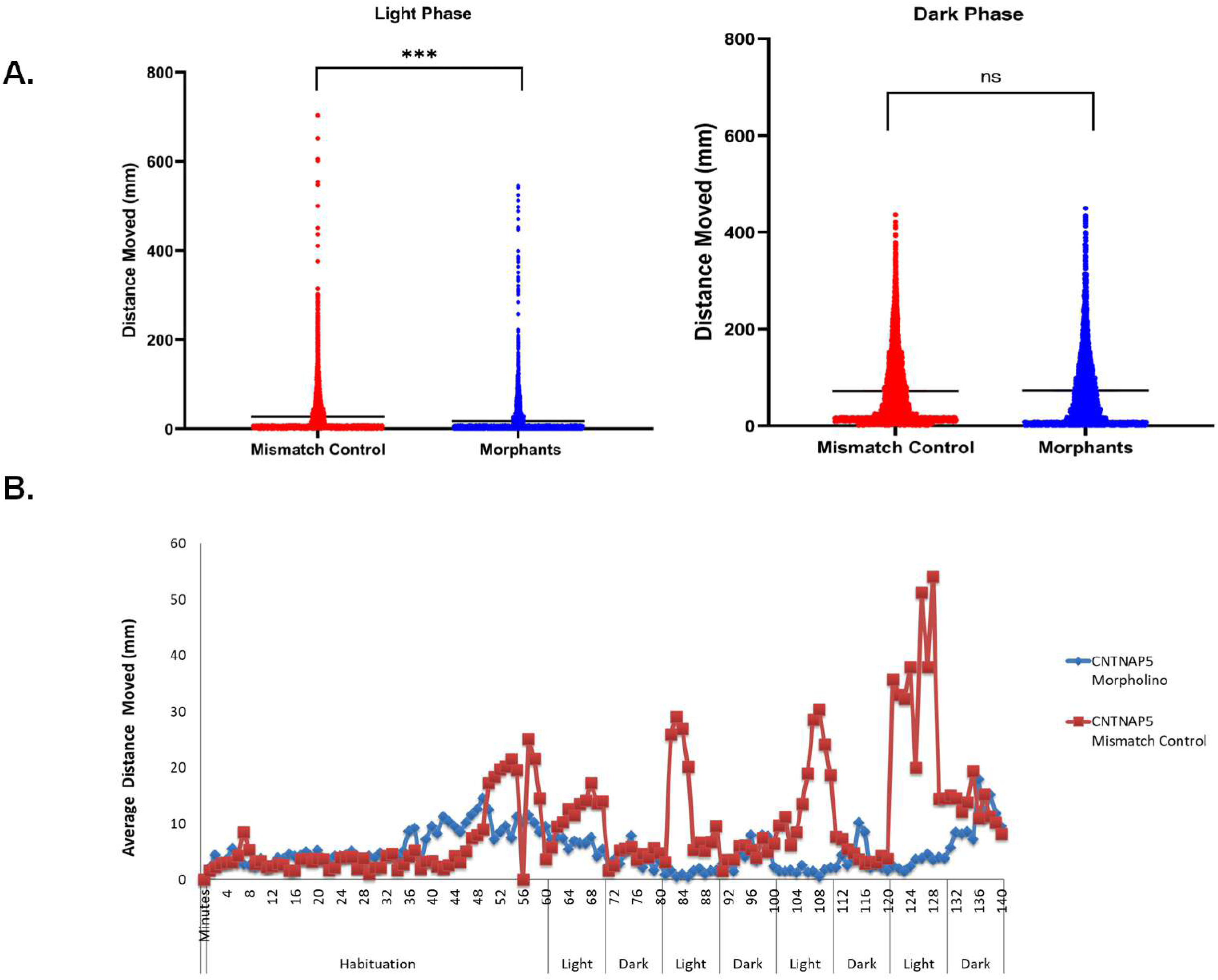
Locomotory movement analysis of zebrafish. **A**. Total distance moved (mm) of 96 hpf *CNTNAP5* morpholino injected zebrafish (blue circles) and 96 hpf *CNTNAP5* mismatch control (red circles) zebrafish for the light (left image) and light off (right image) periods. Left image, (light on) *p*L=L<0.0001, two-sided Mann–Whitney test, *n*L=L24 *CNTNAP5* morpholino injected zebrafish and n=24 *CNTNAP5* mismatch control, Right image (light off), *p*L=L0.5767, bars = mean ± SEM ns not significant, \*\*\*\**p*L<L0.0001. **B**. Average distance moved (mm) of 96 hpf *CNTNAP5* morpholino injected zebrafish (blue circles) and 5dpf *CNTNAP5* mismatch control (red circles) zebrafish exposed to four cycles of 10Lmin light on and 10Lmin light off after 60-minutes habituation period. Each point represents 1Lmin, *n*L=L24 *CNTNAP5* mismatch control zebrafish and *n*L=L24 *CNTNAP5* morpholino injected zebrafish.

## Discussion

Our findings provide compelling evidence implicating *CNTNAP5* as a novel susceptibility gene in glaucomatous neurodegeneration. Integrative analysis of genetic fine-mapping, chromatin architecture, gene regulatory annotations, and tissue-specific expression strongly prioritized *CNTNAP5* as a putative causal gene at the PACG risk locus on chromosome 2. Further, we found that the clustered intronic and downstream variants at the *CNTNAP5* locus exhibiting genome-wide significant association with PACG, overlapped with hotspots of putative cis-regulatory elements. Also, the three-dimensional organization of chromatin within the nucleus plays an important role in controlling gene expression. Regions of high topological associating domain (TAD) interaction frequency can promote contacts between promoters and distal enhancers or silencers to regulate transcription^26^. TAD boundaries are enriched for insulator elements and architectural proteins that may restrict regulatory interactions^27^. Additionally, the local density and activity of cis-regulatory elements (CREs) such as enhancers within a TAD can further modulate expression^28^. Variants disrupting CREs or TAD architecture could thus alter transcriptional programs underlying disease. Our analysis of the Hi-C chromatin conformation data revealed *CNTNAP5* resides within a TAD thus having the capability for modifying/promoting long-range enhancer-promoter interactions. Further, dual luciferase reporter assays provided functional validation confirming allele-specific enhancer activity for rs2553628 residing in *CNTNAP5* region, with the PACG risk allele showing significantly higher activity in the background of constitutive SV40 promoter. Having the strong LD with rs2553628, rs17011429 was found as co-eQTL for schizophrenia^29^.

As per ocular tissue expression database profile, retinal cell types demonstrated marked enrichment for *CNTNAP5* expression compared to other ocular tissues. Furthermore, pathway analysis of *CNTNAP5* co-expressed genes pointed towards involvement in neuronal processes. Furthermore, HumanBase data support the hypothesis that *CNTNAP5* may play a role in retinal disease pathogenesis. The low but measurable confidence score from HumanBase suggests this is a novel association worth further investigation. Additional cross-referencing with the PheWAS catalogue revealed strong genetic correlations of *CNTNAP5* polymorphisms with several neurodevelopmental and neurological disorders. This evidence converges to highlight *CNTNAP5* as a strong functional candidate gene for PACG through its neural-specific expression and role in neurodegeneration. *CNTNAP5*, belongs to the neurexin superfamily of proteins. It plays a crucial role in maintaining the integrity of neural and retinal networks^9^. The *CNTNAP5* gene is a member of the contactin-associated protein (*CNTNAP*) family, which plays a critical role in cell adhesion and signalling. *CNTNAP5* is expressed in the developing and adult nervous system, where it is involved in a variety of processes, including neuronal migration, axon guidance, and synaptic plasticity^30^. The *CNTNAP5* has previously been associated with the posterior cortical atrophy variant of Alzheimers Disease at genome-wide significance^31^, Biopolar disorder^32^, and with schizophrenia^33^. Mutations in the *CNTNAP5* gene have been linked to a number of neurodegenerative diseases, including Autism Spectrum Disorders, Intellectual Disability, and epilepsy^11^.

Using antisense knockdown experiments in zebrafish, we demonstrated cntnap5 knockdown results in ocular phenotypes including reduced eye size, disrupted retinal architecture with discontinuous nuclear layers as observed with reduced NeuroTrace intensity and increased acetylated tubulin levels. Increased acetylated tubulin levels following cntnap5 indicate alterations in microtubule dynamics in cntnap5 morphant retinas possibly leading to cytoskeletal defects and disturbed cytoarchitecture. Acetylation of tubulin is a post-translational modification that alters microtubule stability and intracellular trafficking^34^. Earlier reports suggested that tubulin acetylation restricts axon over branching by reducing microtubule plus-end dynamics in nerves^35^. Defects in microtubule-based transport could disrupt inner retinal laminar organization which manifest as gaps and discontinuous histology. Dysfunctional intracellular trafficking resulting from tubulin hyperacetylation provides a plausible mechanism contributing to the neurodegenerative effects of *CNTNAP5* knockdown. Axonal transport defects are linked to several neurodegenerative diseases^36,37^.

Our findings reveal a significant reduction in NeuroTrace stain intensity in the retinal neurons of cntnap5 morphant zebrafish compared to mismatch control zebrafish. The decreased NeuroTrace stain intensity indicates loss of Nissl substance and suggests the presence of neurodegeneration in the morphant zebrafish retina. The loss of Nissl staining we observed in the neuro-retinal layers of cntnap5 morphants reflects chromatolysis, or disintegration of Nissl bodies, which is a characteristic feature of neurodegeneration^38^. The reduction in NeuroTrace intensity specifically in the outer nuclear layer of the retina indicates that photoreceptor neurons are particularly impacted by cntnap5 knockdown. This aligns with previous studies showing *CNTNAP5* is selectively expressed in vertebrate photoreceptors and suggests an essential role for this protein in maintaining photoreceptor health, likely due to impaired *CNTNAP5* support of neuron structure and function.

Our findings of increased cleaved caspase 3 immunofluorescence in the wholemount cntnap5 knockdown zebrafish provides evidence that deficiency of this gene leads to greater apoptosis during retinal development. The caspase family of cysteine proteases play a central role in programmed cell death pathways^39^. Executioner caspases including caspase 3 are activated by initiator caspases in response to pro-apoptotic signals, ultimately leading to widespread proteolysis and cell death^40^. Increased cleaved caspase 3 staining in the cntnap5 morphant retina indicates enhanced activation of apoptotic pathways resulting in higher levels of programmed cell death. Our observed increase in apoptotic marker caspase 3 suggests cntnap5 plays an important role in retinal cell survival. Cntnap5 deficiency likely makes developing retinal neurons more vulnerable to apoptosis, resulting in thinner retinal layers as observed on histology. Prior studies have similarly implicated dysregulated apoptosis due to inadequate pro-survival signalling in the neurodegenerative effects of glaucoma^41^. Loss of retinal ganglion cells through caspase-mediated apoptosis is a hallmark of glaucomatous pathology^42^. Our finding of increased cleaved caspase 3 parallels reports of enhanced caspase activity in animal models and human glaucoma tissues^43^. Excess apoptosis resulting from cntnap5 deficiency provides a plausible mechanism linking this novel glaucoma risk gene to apoptotic death of retinal neurons. Nonetheless, our cleaved caspase 3 data establish a clear association between *CNTNAP5* knockdown and increased apoptosis in zebrafish larvae.

Finally, our analysis of visually guided behaviour provides functional evidence that cntnap5 knockdown impairs vision in larval zebrafish. Analysis of locomotor activity clearly showed significantly decreased movement during intermittent light periods in cntnap5 morphants compared to controls However, swimming distance in dark conditions was equivalent between groups. This selective reduction in light phase locomotion indicates the cntnap5 deficient larvae have impaired visual function despite intact neuromuscular capacity. Similar light/dark dependent behavioural defects have been reported in zebrafish models of inherited blindness^44^. Visually-guided locomotion requires integration and transmission of visual cues to motor circuits^45^. Decreased photoperiod movement in cntnap5 morphants reflects an inability to respond appropriately to light onset due to deficiencies in retinal processing. Loss of cntnap5 expression likely impairs retinal circuit development and function, manifesting as vision-dependent locomotor deficits. Further electrophysiological analysis of visual responses and circuit connectivity in cntnap5 morphants will provide greater mechanistic detail into how cntnap5 loss impacts retinal signalling pathways leading to functional vision loss. However, our locomotor activity data clearly demonstrates cntnap5 plays a critical role in the proper development of visual behaviour and function in zebrafish larvae. Our findings are consistent with the morphological and histological retinal abnormalities resulting from cntnap5 knockdown underlying vision loss.

This study has some limitations that should be considered when interpreting the identification and prioritization of *CNTNAP5* as a PACG risk gene. First, eQTL data specific to ocular tissues is not available in GTEx or other databases, preventing assessment of whether the PACG-associated variants act as eQTLs on CNTNAP5 expression (GTEx Consortium). Additionally, the lack of ocular eQTL data precluded colocalization analysis to determine if the PACG and eQTL signals are driven by the same underlying causal variant(s)^46^. While enrichment of regulatory annotations provides supportive functional evidence, eQTL data from disease-relevant tissues would further strengthen *CNTNAP5* as the likely causal gene. Next, while examination of Hi-C data suggested *CNTNAP5* resides in a TAD, 3C-based sequencing was not performed to directly measure chromatin looping interactions with the risk variants^47^. Further 3C analysis will be needed to validate direct physical contacts between the PACG-associated variants and *CNTNAP5* promoters or enhancers, which is beyond scope of this study. Finally, although conditional analysis pointed to *CNTNAP5* over other nearby genes, additional functional experiments are required to definitively confirm its role in PACG pathogenesis. However, taken together, the genetic fine-mapping, bioinformatics annotation, and preliminary zebrafish knockdown results provide reasonable prioritization of *CNTNAP5* as a novel PACG risk gene despite these limitations. Future studies overcoming these caveats will provide greater certainty into the regulatory mechanisms and causal role of *CNTNAP5* variants in PACG aetiology.

In conclusion, our integrated genomic, expression, and functional analyses provide compelling evidence for *CNTNAP5* as a novel PACG risk gene conferring susceptibility through altered regulation in retinal cell types. The effects of cntnap5 knockdown in zebrafish further validate its role in neurodegeneration. Given its neural-specificity and involvement in cytoskeletal dynamics, *CNTNAP5* represents a promising target for future investigation and therapeutic development for PACG and related neurodegenerative conditions. Further studies should explore how modulation of cntnap5 expression influences intracellular trafficking and survival signalling in retinal ganglion cells. Elucidating the precise molecular mechanisms through which dysregulated *CNTNAP5* contributes to PACG pathogenesis will yield crucial insights into disease aetiology while nominating strategies for targeted intervention.

## Acknowledgements

This work was supported by NIBMG intramural funds to Dr. Moulinath Acharya and Indian Council of Medical Research (ICMR) senior research fellowship of Sudipta Chakraborty (No.3/1/3(8)/OPH/2020-NCD-II).

## Author Contribution

**SC** performed all the statistical, bioinformatic functional and behavioural analysis, conceived the study, drafted and edited the original manuscript. **JS** performed luciferease assay and western blot. **SSR** performed zebrafish tissue processing and maintenance. **SMi** performed immunofluorescence and confocal imaging. **SB** performed whole mount immunofluorescence **SD** performed immunofluorescence. **SS** performed bioinformatic analysis **SMa** performed confocal imaging **SB** supervised all the statistical and bioinformatics analyses, helped in conceiving and designing the study, reviewing and editing of the manuscript draft. **MM** supervised all the functional analyses, helped in conceiving and designing the study, reviewing and editing of the manuscript draft. **MA** conceived the study, supervised all the analyses and interpretation of results and drafted the manuscript. All authors read and approved the final version of the manuscript for publication.

