## Supplementary information for "Post-GWAS functional analyses of *CNTNAP5* suggests its role in glaucomatous neurodegeneration"

| Morpholino | Sequence |
| --- | --- |
| cntnap5a | TTAAGATTTGGTCTTGTTGGCATCT |
| cntnap5a_5_base_mismatch | TTAAcATTTcGTgTTcTTGcCATCT |
| cntnap5b | GGCCGAGCGAAGATATTCCATGTTC |
| cntnap5b_5_base_mismatch | GGCCcAcCGAAcATATTCgATcTTC |

**Table S1**: Morpholino sequence of cntnap5a, cntnap5b and their respective mismatch 5 base mismatch control.

| Chromosome | SNP | RegulomeDB Rank |
| --- | --- | --- |
| 2 | rs17011420 | 3a |
| 2 | rs17724018 | 4 |
| 2 | rs2115890 | 4 |
| 2 | rs17011429 | 5 |
| 2 | rs2901264 | 5 |
| 2 | rs1430263 | 5 |
| 2 | rs779979 | 5 |
| 2 | rs17011399 | 6 |
| 2 | rs780010 | 6 |
| 2 | rs733112 | 6 |
| 2 | rs2553625 | 7 |

**Table S2:** RegulomeDB score of 13 SNPs of *CNTNAP5*

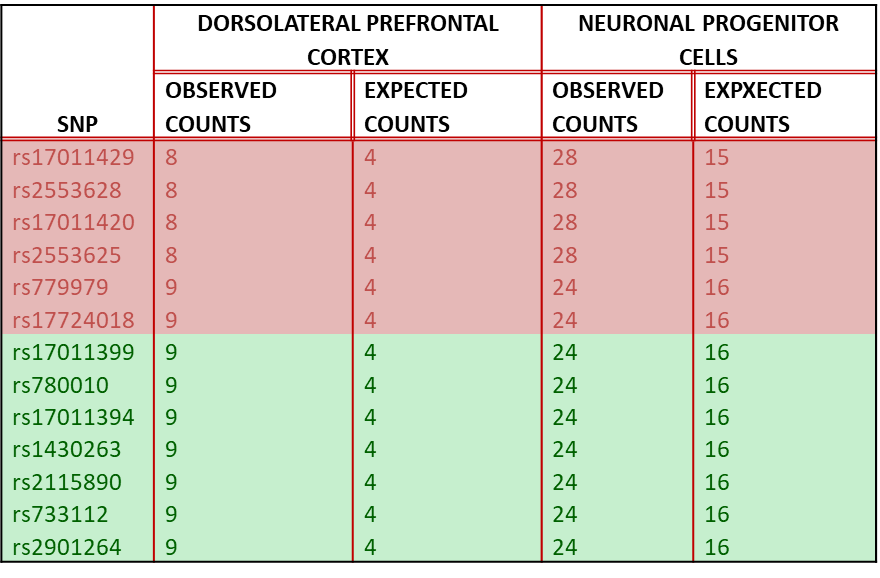

**Table S3**: Observed and expected counts of the intronic SNPs of CNTNAP5 found from HUGIn. These SNPs that were distributed in two LD blocks (the SNPs in pink belong to one LD block and the ones in green belonged to the other LD block) and showed similar observed and expected read counts.

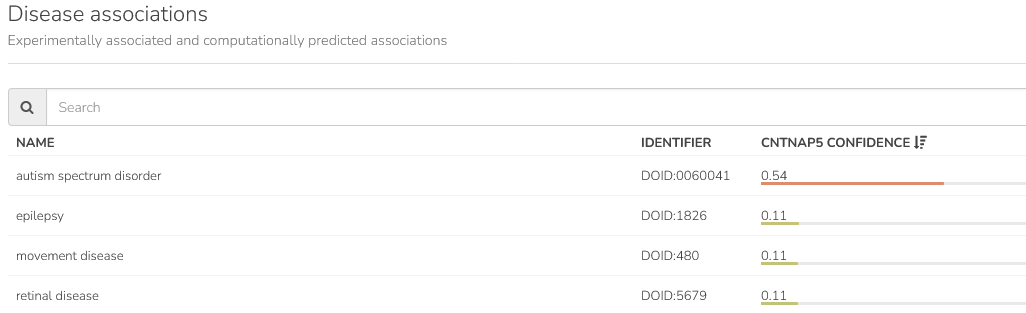

**Table S4**: HumanBase portal showed *CNTNAP5* as the highest confidence score for autism spectrum disorder. The gene also showed low but measurable confidence scores for epilepsy and retinal disease. Other neurodegenerative diseases like Parkinson's and Alzheimer's showed very low confidence scores.

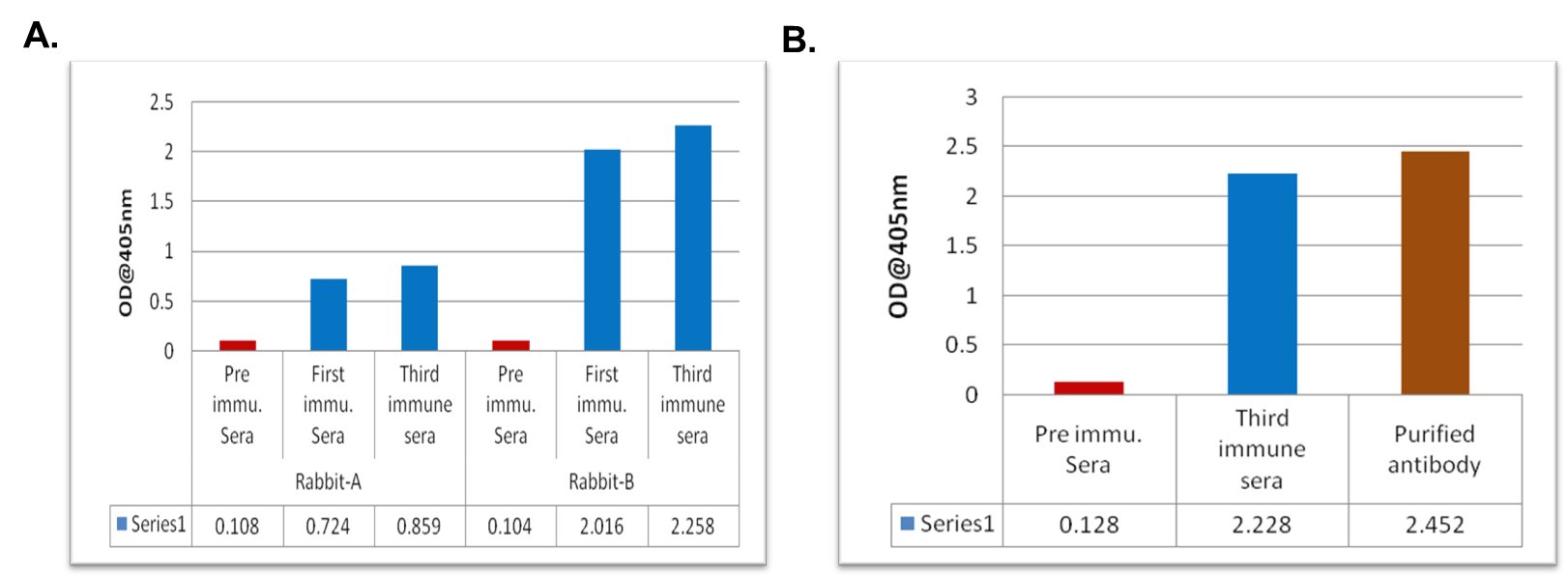

**Figure S1 A:** First and Third immune sera of Rabbit A and B tested against the antigen coated at 200ng/well and at 1:5000 dilutions of primary Ab. Pre-immune sera were used as control in place of primary antibody. Plates read after 15 min of enzyme substrate reaction and the absorbance were measured at 405nm. **B.** Immune sera (at 1:5000) and Purified antibody (at 200ng/well) tested against the antigen coated at 200ng/well obtained a value of 2.228 and 2.452 respectively. Pre-immune sera were used as control in place of primary antibody. Plates read after 15 min of enzyme substrate reaction and the absorbance were measured at 405nm.

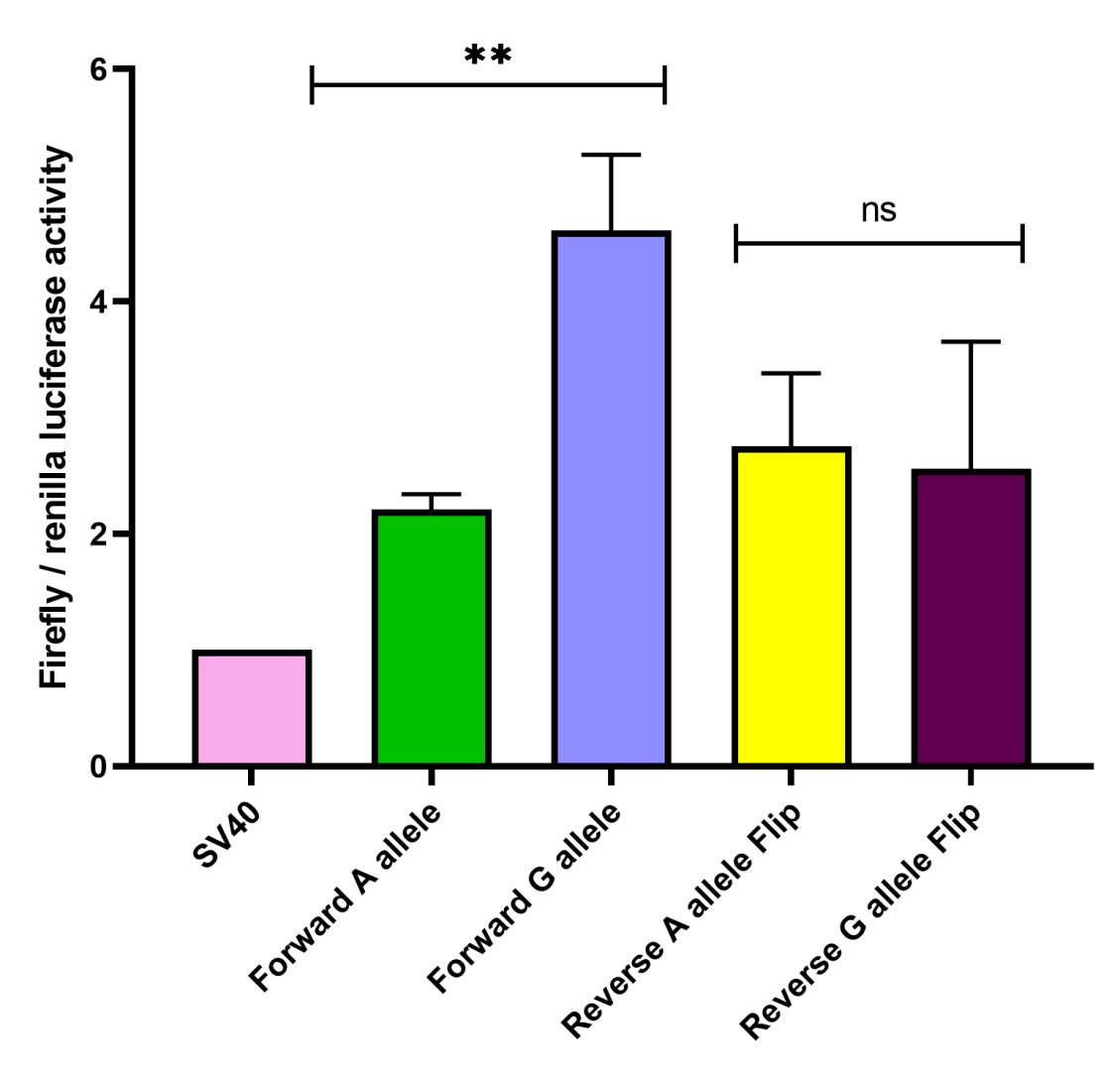

**Figure S2:** The data was drawn for a total of 3 biological replicates considering triplicate technical replicates for each biological experiments representing three different plasmid preparations (pGL3 SV40 minimal promoter vector) transfected in to HEK293T cells grown in 96-well plates. The bar plots the mean value of all the 3 experiments, while error bars are depicting SD. Two tailed student t-test for independent means was used for calculating statistical significance; *p < 0.05, **p < 0.01; ***p < 0.001, ns-= not significant. The luciferase assay shows G allele of rs2553628(CNTNAP5) has a statistically significant higher firefly / renilla luciferase activity than the A allele of rs2553628 (*CNTNAP5*) for both forward orientation and reverse orientation. P-value: 0.0034 (Forward set) P-value: 0.8041 (Reverse set).

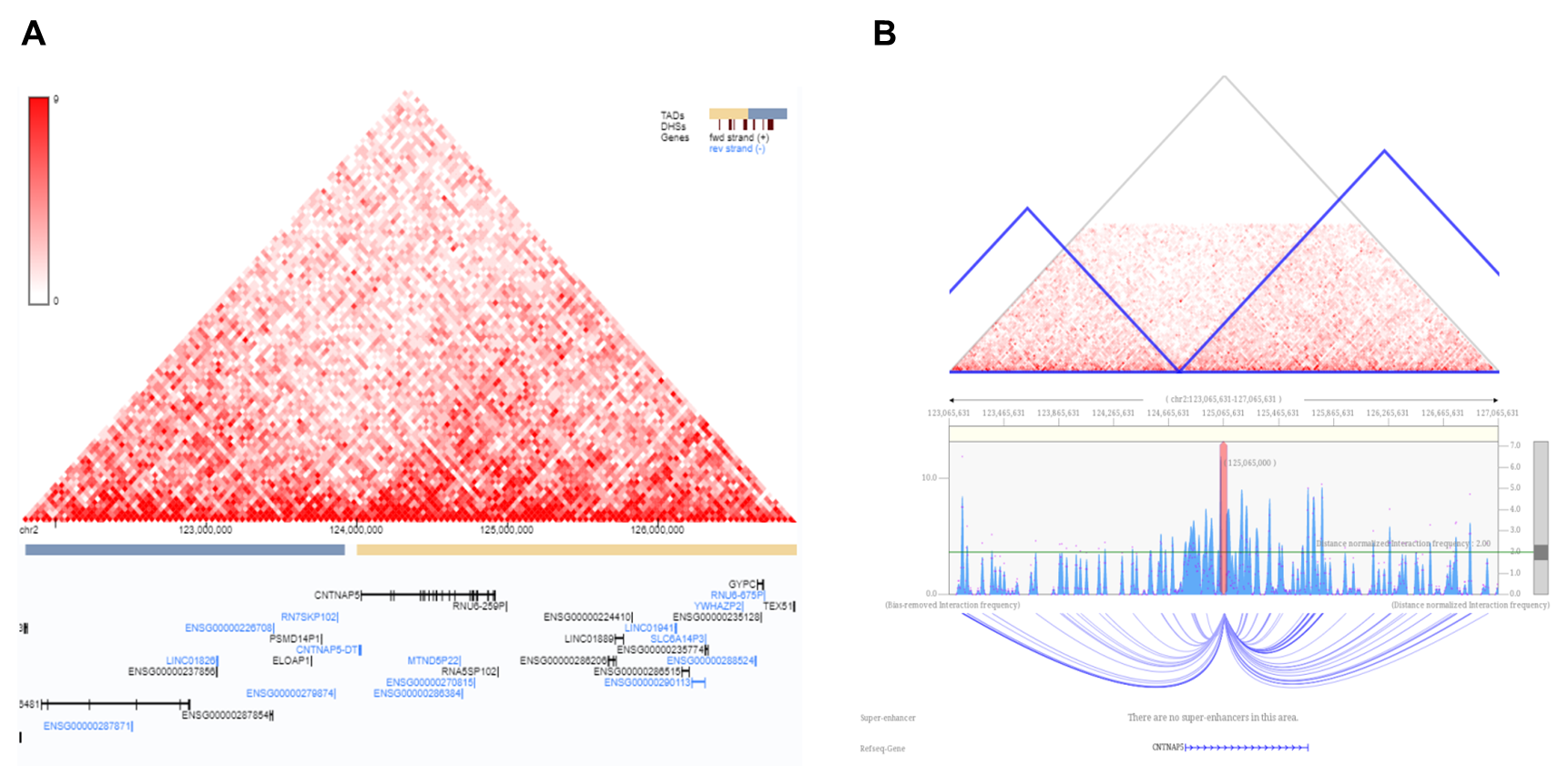

**Figure S3**: **A**. The Hi-C heatmaps clearly show strong TAD correlation around genomic region chr2:125,083,095-125,084,384 of *CNTNAP5*. **B.** Visualization of significant long interaction arcs centred on genomic coordinates chr2:125,083,095-125,084,384 located at the *CNTNAP5* intron. From the top, normalized Hi-C contact map with TAD annotations (in blue triangles), identified chromatin interactions and the RefSeq genes. The blue bar graph represents the bias-removed interaction frequencies and the magenta dots represent the distance normalized interaction frequencies. The blue bars above the green threshold line are the interactions which are 2-fold greater than the background listed ones.

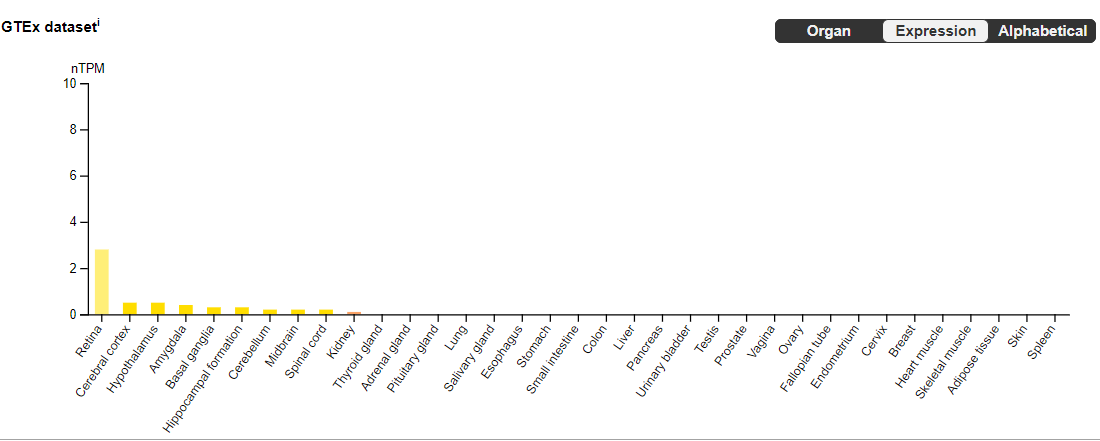

**Figure S4**: Expression of cntnap5 is restricted only in retina and neural tissues (*Source: Human Protein Atlas*).

normalized reads

|  | mean |
| --- | --- |
| Healthy conjunctiva | 0 |
| Cornea | 0.55 |
| Lid | 0 |
| Healthy lacrimal gland | 3.825 |
| Optic nerve | 7.56667 |
| Healthy retina periphery | 5.2 |
| Healthy retina central | 34.925 |
| Healthy choroid RPE | 0 |
| Retinal microglia | 0.121429 |

**Figure S5:** Expression profile of cntnap5 in the different eye tissue (*Source: Human Eye Transcriptome Atlas*)

coex-z

|  | coex z |
| --- | --- |
| GABRG2 | 8.5 |
| SLC17A6 | 7.9 |
| KCNH5 | 7.5 |
| ELAVL4 | 7.3 |
| CSMD3 | 7.2 |
| PAK5 | 7.1 |
| CLVS2 | 7.1 |
| OLFM3 | 7 |
| NXPH1 | 6.9 |
| GRIA2 | 6.9 |
| CA10 | 6.8 |
| CNTNAP2 | 6.8 |
| DSCAM | 6.8 |
| DPP10 | 6.3 |
| CDH8 | 5.8 |
| DCC | 5.8 |
| OPRK1 | 5.2 |
| ADAMTS18 | 4.4 |
| KCNMB2 | 3.7 |

**Figure S6**: Co-expressed gene of *CNTNAP5*. Co-expression values as coex-z (a coex z values greater than 3 is significant it implies to an FDR of 0.1% also ruling out that they are randomly co-expressed.

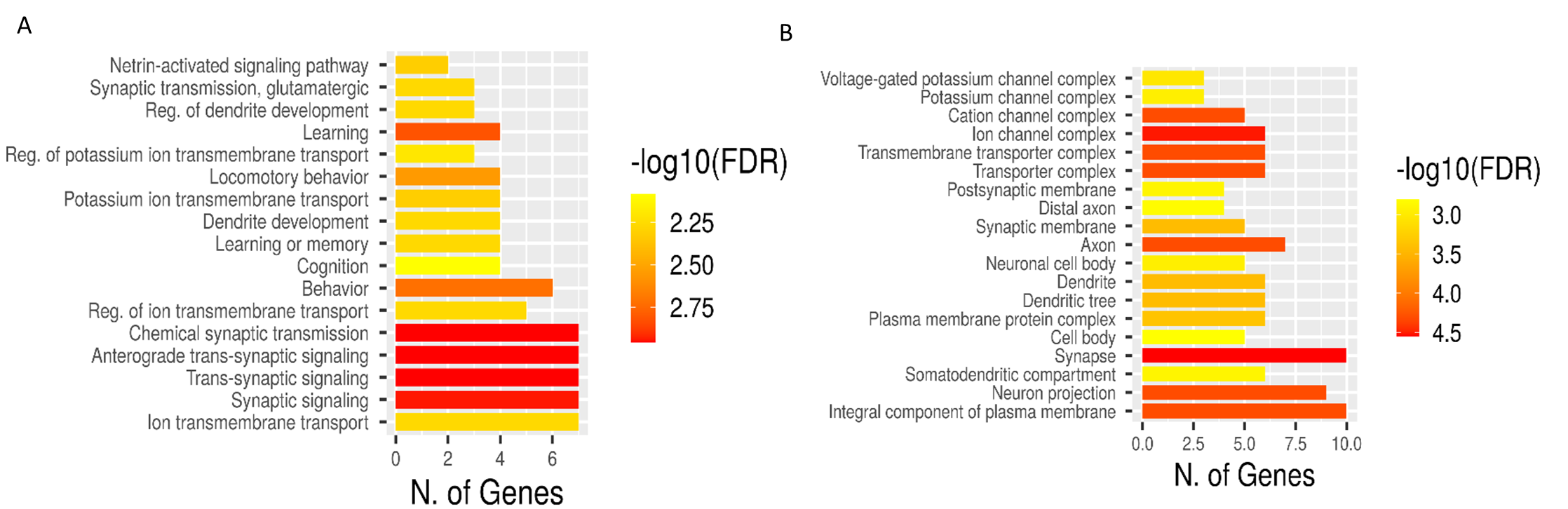

**Figure S7:** The gene enrichment analysis of the co-expressed genes of *CNTNAP5* from the following databases-**A**. GO Biological Process **B**. GO Cellular Components.

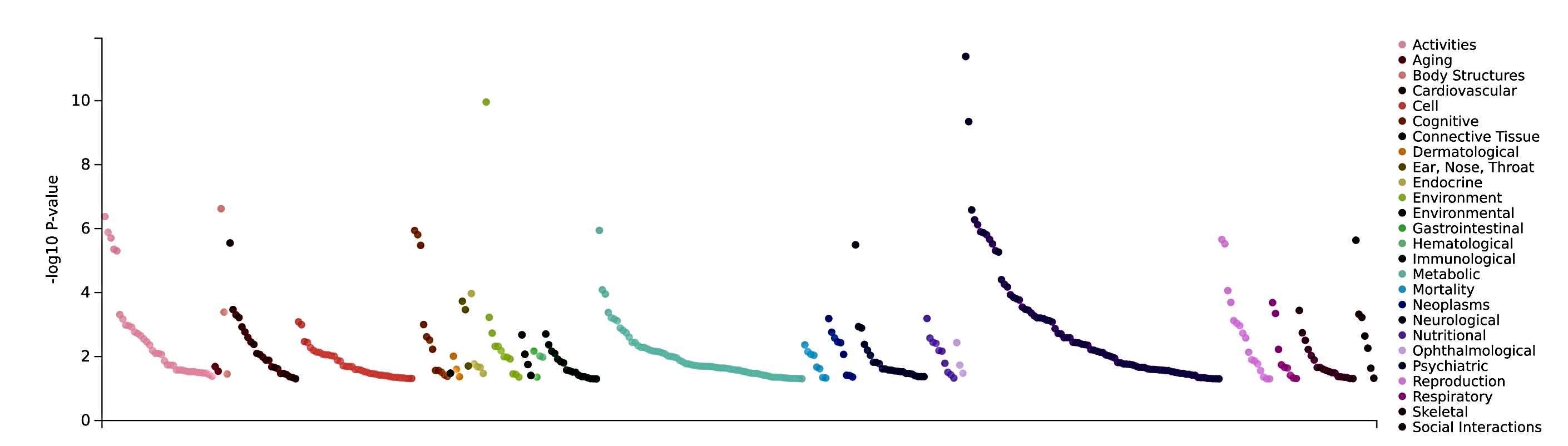

**Figure S8**: The disease annotations of *CNTNAP5* as per the data of PheWAS catalog.

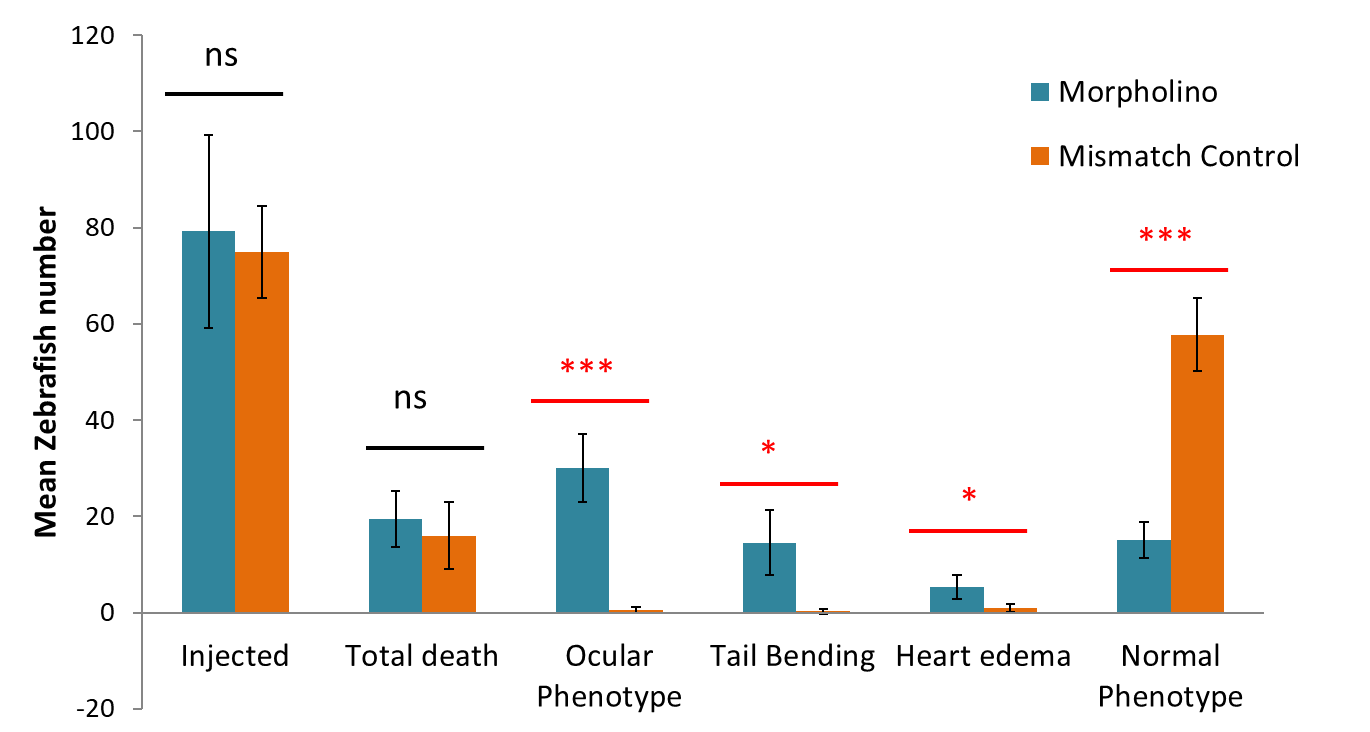

**Figure S9**: The bar plots the mean value of all the 8 experiment sets, while error bars are depicting SD. Two tailed student *t*-test for independent means was used for calculating statistical significance; *p < 0.05, **p < 0.01; ***p < 0.001, ns-= not significant.

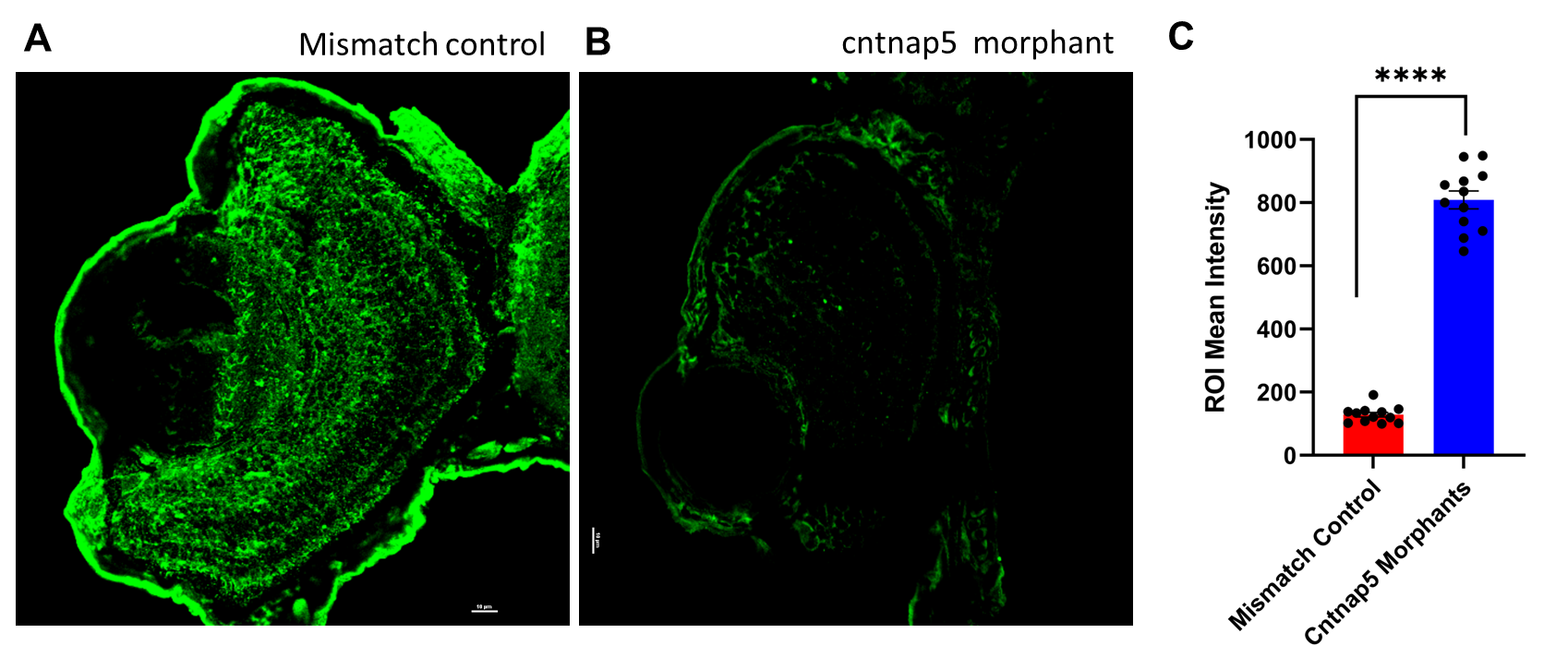

**Figure S10**: Representative confocal images of cntnap5 expression of eye tissues from mismatch control fish **(A)** and cntnap5 morphant **(B)** zebrafish at 96 hpf. **C.** Comparative analysis of mean intensity of eye for the both groups, bars = mean ± SE, ns not significant, *****p* < 0.005.

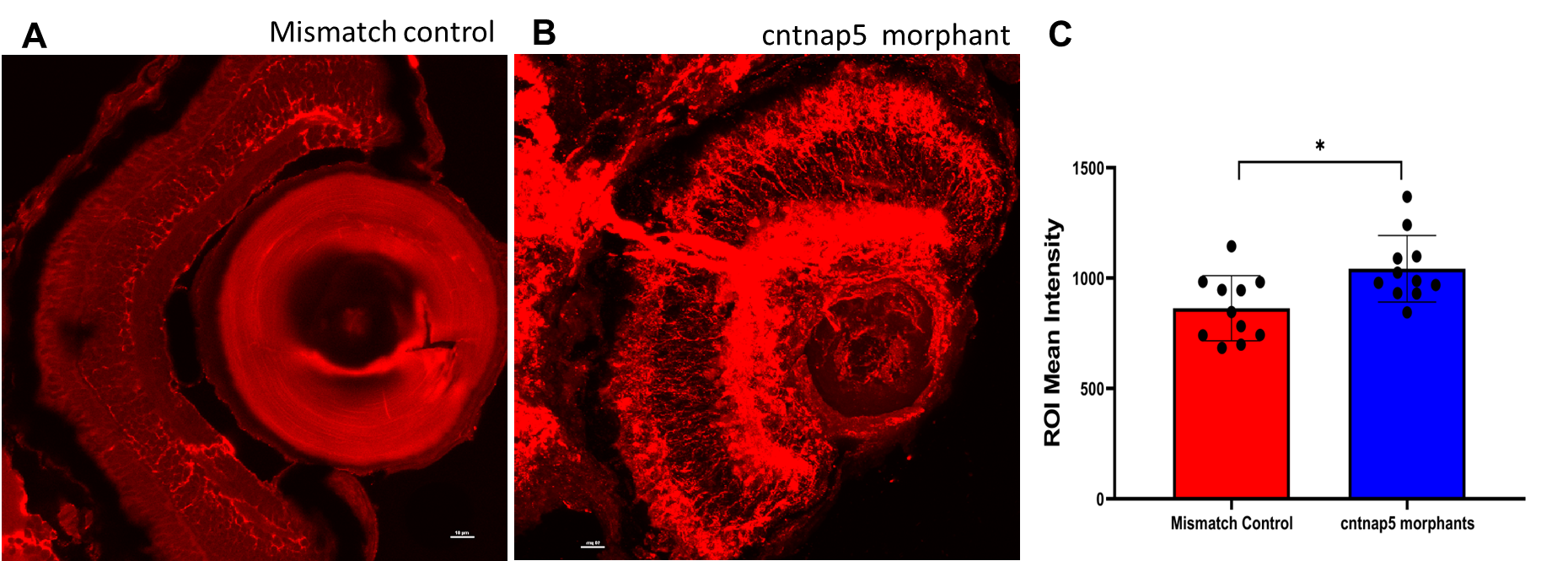

**Figure S11**: Representative confocal images of acetylated tubulin expression of eye tissues from mismatch control fish **(A)** and cntnap5 morphant **(B)** zebrafish at 96 hpf. **C.** Comparative analysis of mean intensity of eye for the both groups, bars = mean ± SE, ns not significant, ****p* < 0.005.

**Video S1**: Locomotory movement video of zebrafish. The distance moved of 5dpf *CNTNAP5* mismatch control (upper 3 wells) zebrafish and 96hpf *CNTNAP5* morpholino injected zebrafish (lower 3 wells) exposed to four cycles of 10 min light on and 10 min light off period.
